# Chromatin-Assisted Targeting Enables Precise DNA Methylation Editing in Plants

**DOI:** 10.64898/2026.08.25.747117

**Authors:** Yan He, Ming Wang, Tyler J. Buckley, Brandon A. Boone, Erika Li, Janice Yerin Shin, Nicholas Alvarado, Bohao Xu, Anh Nguyen, Shuya Wang, Yuxing Zhou, Suhua Feng, Steven E. Jacobsen

## Abstract

Precise installation of DNA methylation at selected loci offers a powerful strategy for regulating gene expression without altering DNA sequence, but existing plant epigenome editors are constrained by limited efficiency, locus dependence, and genome-wide off-target methylation. Here, we developed SunTag-MQ1v variants incorporating TRBIP1, which promotes removal of the antagonistic H3K4me3 mark, and CHLAMY, an oligomerizing alpha crystalline domain protein from *Chlamydomonas reinhardtii*. TRBIP1 enhanced methylation and silencing at the *Arabidopsis FWA* promoter but caused widespread off-target methylation and severe developmental defects. Adding CHLAMY produced SunTag-CHLAMY-TRBIP1-MQ1v (designated as SunTag-NOVA), which successfully overcame the lethality and widespread off-target effects associated with direct TRBIP1-MQ1v fusions. We demonstrate that CHLAMY drives higher-order oligomerization of the editing complex, which enhances target specificity and mitigates off-target accumulation. SunTag-NOVA robustly installed DNA methylation and repressed transcription at the endogenous *FWA, FT* and *TMM* genes with minimal genome-wide off-target consequences. These results show that combining local chromatin modification with controlled effector assembly can improve targeted DNA methylation, and establish SunTag-NOVA as a specific epigenome-editing platform for plants.

## Introduction

DNA methylation is a fundamental epigenetic mark that regulates gene expression, maintains genome stability, and safeguards normal development^1,2^. In plants, *de novo* methylation is primarily catalyzed by the RNA-directed DNA methylation (RdDM) pathway, in which 24-nt small interfering RNAs (siRNAs) guide the methyltransferase DRM2 to deposit methylation in all sequence contexts (CG, CHG, and CHH)^1,2^. Once established, DNA methylation is faithfully propagated across cell divisions by distinct maintenance mechanisms: the CG methyltransferase MET1 preserves symmetric CG methylation, while CMT3 maintains CHG methylation and CMT2 reinforces CHH methylation at heterochromatin^3^. The development of targeted DNA methylation tools is essential for precisely modifying epigenetic states at defined genomic loci without globally perturbing the epigenome.

CRISPR/dCas9 (dead Cas9)-based approaches are widely employed to achieve targeted DNA methylation and locus-specific gene silencing across plant and mammalian systems^4-6^. In *Arabidopsis*, CRISPR/dCas9-based systems employing the catalytic domain of *Nicotiana tabacum* Domains Rearranged Methyltransferase (NtDRMcd) or a CG-specific bacterial methyltransferase variant (MQ1v) have been well documented for targeted DNA methylation. However, the dCas9-NtDRMcd system induces substantial genome-wide off-target hypermethylation, whereas the dCas9-based MQ1v system shows reduced off-target effects but has been effective only at the *FWA* locus^4, 6^. To overcome these limitations, we aimed to engineer a dCas9-based MQ1v system that minimizes off-target methylation and enhances its general applicability.

We previously showed that removal of histone H3 lysine 4 trimethylation (H3K4me3) by the TRB INTERACTING PROTEIN1 (TRBIP1), which recruits the H3K4me3 demethylase JMJ14 to target loci, can enhance targeted DNA methylation^7^. In other work, α-crystalline domain (ACD) proteins have been shown to stimulate the multimerization of the methyl DNA binding proteins MBD5 and MBD6 in their normal role in regulating expression at DNA methylation sites in *Arabidopsis*, suggesting a critical role for ACD proteins in promoting protein assembly^8,9^. Building on these findings, we combined TRBIP1-mediated H3K4me3 demethylation and ACD protein fusion with a CRISPR/dCas9-based MQ1v system to develop a novel epigenome-editing tool with high specificity, efficiency, and general applicability in *Arabidopsis*, with potential for deployment across diverse plant species.

## Results

### Efficiency enhancement of TRBIP1 in a SunTag-MQ1v system

The SunTag system uses a dCas9 protein fused to 10x GCN4 peptide repeats, which is directed to specific genomic loci by guide RNAs (gRNAs). These GCN4 repeats are then recognized by a GFP-tagged single-chain variable fragment (scFv-GFP) fused to an effector protein, thereby recruiting the effector to the targeted genomic site^10^. We previously established a SunTag-MQ1v (MQ1v is a variant of the CG-specific bacterial methyltransferase SssI) system to add DNA methylation to specific loci in *Arabidopsis* genome^4^. Due to the limited efficiency of this SunTag-MQ1v system, we pursued engineering approaches aimed at improving its overall capability. Recently, we described a small coiled coil domain protein called TRB INTERACTING PROTEIN1 (TRBIP1) that potently removes H3K4me3 to silence gene expression in Arabidopsis^7^. Motivated by these findings, we examined whether the direct fusion of TRBIP1 with MQ1v in the SunTag system could improve its functionality.

In *Arabidopsis, FLOWERING WAGENINGEN* (*FWA*) is frequently employed as a methylation reporter, as its promoter is typically heavily methylated and transcriptionally silenced in wild-type (WT) plants. However, in the *fwa* epiallele mutant, methylation at the *FWA* promoter is permanently lost, leading to *FWA* overexpression and a characteristic late-flowering phenotype^6, 11, 12^. We introduced the SunTag-TRBIP1-MQ1v_*FWA*-gRNA construct (with two gRNAs directed to *FWA* promoter) into the *fwa* mutant background. Compared with SunTag-MQ1v_*FWA*-gRNA, the SunTag-TRBIP1-MQ1v_*FWA*-gRNA construct triggered a much stronger early-flowering phenotype in T1 plants. However, these early-flowering individuals also exhibited severe growth defects, with most plants dying shortly after bolting and failing to produce seeds (Fig. 1A,B and Supplementary Fig. 1A). We examined DNA methylation levels at the *FWA* promoter and found that SunTag-TRBIP1-MQ1v_*FWA*-gRNA induced substantially higher methylation deposition compared with SunTag-MQ1v_*FWA*-gRNA (Fig. 1C,D). As expected, *FWA* expression was efficiently reduced to WT levels in the *fwa* background expressing SunTag-TRBIP1-MQ1v_*FWA*-gRNA (Fig. 1E), consistent with its strong early-flowering phenotype and enhanced promoter methylation relative to SunTag-MQ1v_*FWA*-gRNA. Whole-genome bisulfite sequencing (WGBS) further revealed that SunTag-TRBIP1-MQ1v_*FWA*-gRNA caused very extensive off-target methylation across the genome (up to 50% CG methylation), as well as over 70% methylation of the chloroplast genome (Fig. 1F,G and Supplementary Fig. 1B,C). Such widespread hypermethylation likely contributes to the lethality and seed-set defects observed in the SunTag-TRBIP1-MQ1v_*FWA*-gRNA lines. Taken together, our results indicate that although TRBIP1 greatly enhances the methylation-inducing activity of SunTag-MQ1v, the accompanying off-target effects substantially limit the utility of SunTag-TRBIP1-MQ1v as an epigenetic editing tool.

**Fig. 1.**
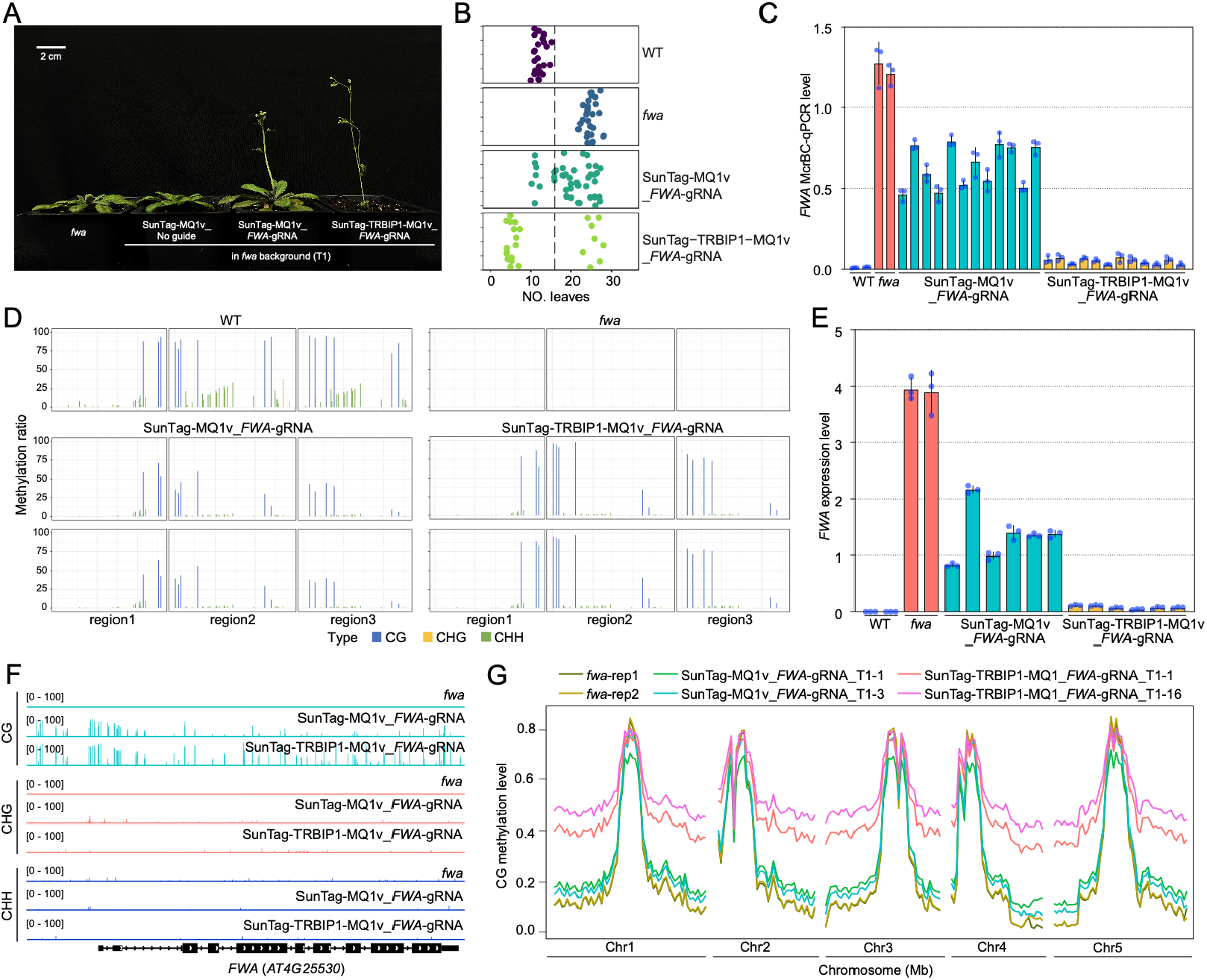
TRBIP1 enhances the DNA methylation efficiency of the SunTag-MQ1v system. (*A*) Phenotypic comparison of T1 plants expressing SunTag-MQ1v_*FWA*-gRNA or SunTag-TRBIP1-MQ1v_*FWA*-gRNA in the *fwa* background. (*B*) Flowering time quantified by counting leaf number prior to flowering in SunTag-MQ1v_*FWA*-gRNA or SunTag-TRBIP1-MQ1v_*FWA*-gRNA T1 plants. (*C*) DNA methylation at the *FWA* promoter in SunTag-MQ1v_*FWA*-gRNA or SunTag-TRBIP1-MQ1v_*FWA*-gRNA T1 plants measured by McrBC-qPCR; lower values indicate higher methylation levels. (*D*) BS-PCR sequencing showing CG, CHG, and CHH methylation patterns at the *FWA* promoter in WT, *fwa*, two T1 SunTag-MQ1v_*FWA*-gRNA lines, and two T1 SunTag-TRBIP1-MQ1v_*FWA*-gRNA lines in the *fwa* background. (*E*) *FWA* expression levels in WT, *fwa*, SunTag-MQ1v_*FWA*-gRNA and SunTag-TRBIP1-MQ1v_*FWA*-gRNA T1 lines in *fwa*. (*F*) Genome browser snapshot illustrating DNA methylation profiles across *FWA* promoter in *fwa*, SunTag-MQ1v_*FWA*-gRNA, and SunTag-TRBIP1-MQ1v_*FWA*-gRNA T1 lines. (*G*) Comparison of whole-genome CG methylation levels between two SunTag-MQ1v_*FWA*-gRNA and SunTag-TRBIP1-MQ1v_*FWA*-gRNA T1 lines in the *fwa* background. For (*C*) and (*E*), data are presented as mean ± SD (*n* = 3).

### Engineering diverse ACD proteins for the SunTag-TRBIP1-MQ1v system

Despite the significant improvement in SunTag-MQ1v activity resulting from the direct fusion of TRBIP1 to MQ1v (Fig. 1), two major challenges remain: the associated lethality and widespread off-target methylation across the genome. The α-crystallin domain (ACD), a defining feature of the small heat shock protein (sHSP) family, spans roughly 90 amino acids and is flanked by variable N- and C-terminal regions^13^. The ACD assembles into dimeric building blocks, while the flanking terminal domains guide the formation of higher-order oligomers^13^. Our recent study found that α-crystalline domain (ACD) protein ACD15 and ACD21 drive multimerization of methyl-CpG binding domain (MBD) proteins MBD5 and MBD6 to silence target genes and transposable elements (TEs) in Arabidopsis^8^. We sought to use the multimerizing properties of ACD proteins to aggregate excess TRBIP1-MQ1 effector proteins to reduce off target DNA methylation. We therefore searched for ACD15/21 homologs across the kingdoms of life, and fused selected ACDs to the *FWA*-targeting SunTag system. (SunTag-ACD) and tested whether these non-Arabidopsis ACD proteins could support aggregation. Based on the intensity of GFP aggregation foci observed by live confocal imaging of Arabidopsis root cells, we classified the tested ACDs into three categories: strong foci, weak foci, and no foci (Fig. 2A). To evaluate whether incorporating these ACDs into the SunTag-TRBIP1-MQ1v system could alleviate lethality and off-target methylation in Arabidopsis, we selected five representative candidates and fused them to TRBIP1-MQ1v within the SunTag framework (SunTag-ACD-TRBIP1-MQ1v). *Chlamydomonas reinhardtii* (CHLRE_07g318800v5, hereafter referred to as CHLAMY), *Zea mays* (ZmHSP18), and *Homo sapiens* (HsHSPB1) were in the strong-foci group. HsHSPB4 fell into the weak-foci group, whereas HsHSPB8 was classified in the no-foci group. To reduce post-transcriptional gene silencing and reduce variability between transgenic lines, we introduced the different *FWA*-targeting SunTag-ACD-TRBIP1-MQ1v constructs into the *fwa rdr6* double mutant^7^. Among the tested ACD proteins, only CHLAMY both efficiently rescued the lethality phenotype and preserved the robust early-flowering phenotype induced by SunTag-TRBIP1-MQ1v in the *fwa rdr6* background (Fig. 2B). We therefore selected CHLAMY as the candidate for further analysis.

**Fig. 2.**
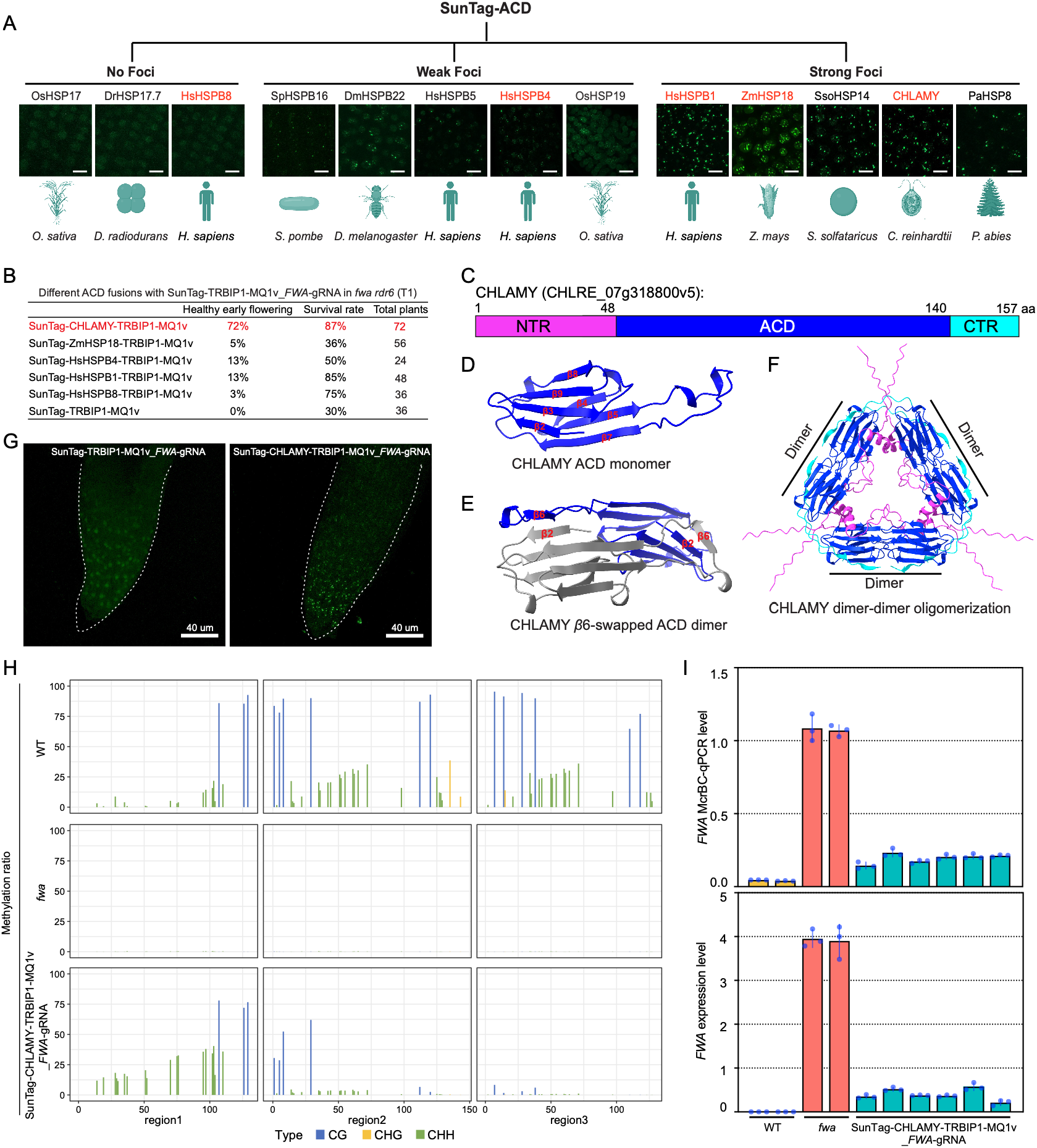
CHLAMY is effective for SunTag-TRBIP1-MQ1v engineering. (*A*) Representative images of SunTag-ACD scFv-GFP localization in root tips using ACDs from different species. ACDs shown in red font represent the candidates selected for incorporation into the SunTag-TRBIP1-MQ1v system. Scale bar = 10 µm. (*B*) Phenotypes of T1 plants expressing different ACD fusions with SunTag-TRBIP1-MQ1v in the *fwa rdr6* background. (*C*) Schematic representation of the CHLAMY domain architecture. NTR, N-terminal region; ACD, alpha-crystallin domain; CTR, C-terminal region. (*D*) AlphaFold-predicted *β*-sandwich structure of the CHLAMY ACD. (*E*) Structures of the *β*6-swapped CHLAMY ACD dimer, with the two monomers shown in gray and blue. (*F*) Model illustrating how CHLAMY NTR and CTR may mediate dimer-dimer oligomerization; NTR shown in magenta and CTR in cyan. (*G*) Root tip svGFP signals in T1 lines expressing SunTag-TRBIP1-MQ1v or SunTag-CHLAMY-TRBIP1-MQ1v in the f*wa* background. (*H*) BS-PCR analysis of CG, CHG, and CHH methylation at the *FWA* promoter in WT, *fwa*, and SunTag-CHLAMY-TRBIP1-MQ1v_*FWA*-gRNA T1 plants in the *fwa* background. (*I*) *FWA* promoter methylation (McrBC-qPCR) and *FWA* expression (qRT-PCR) levels in T1 SunTag-CHLAMY-TRBIP1-MQ1v_*FWA*-gRNA plants. Data represent mean ± SD (*n* = 3).

The primary structure of CHLAMY consists of a conserved, folded ACD core flanked by a flexible N-terminal region (NTR) and a short C-terminal region (CTR) (Fig. 2C). AlphaFold3 predicts that the ACD monomer adopts a *β*-sandwich architecture composed of two antiparallel *β*-sheets containing three and four *β* strands, respectively (Fig. 2D). In addition, the ACD forms dimers through reciprocal insertion of the newly formed *β*6 strand into the *β*-sandwich of the partner monomer, and this interaction was validated using yeast two-hybrid assays (Fig. 2E and Supplementary Fig. 2). The NTR and CTR are typically involved in assembling isolated ACD dimers into higher-order oligomers^14,15^. Consistent with this, CHLAMY dimers are predicted to cross-link through their NTR and CTR to generate larger oligomeric structures (Fig. 2F). We further assessed CHLAMY oligomerization by examining GFP signal aggregation in T1 plants expressing SunTag-CHLAMY-TRBIP1-MQ1v_*FWA*-gRNA. Discrete GFP foci were observed in these plants, whereas GFP fluorescence remained diffuse in SunTag-TRBIP1-MQ1v_*FWA*-gRNA plants lacking CHLAMY (Fig. 2G). As expected from the strong early flowering phenotype, in the *fwa* background, SunTag-CHLAMY-TRBIP1-MQ1v_*FWA*-gRNA efficiently methylated the *FWA* promoter, leading to strong repression of *FWA* expression (Fig. 2H,I). Together, these results indicate that CHLAMY mediates higher-order oligomerization of the SunTag-CHLAMY-TRBIP1-MQ1v complex in Arabidopsis, and that this oligomerization enhances specific methylation targeting and silencing of *FWA*. Given the strong performance of SunTag-CHLAMY-TRBIP1-MQ1v in directing DNA methylation at *FWA* locus (Fig. 2), with minimal ancillary morphological defects, we hereafter refer to this construct as SunTag-NOVA.

### The addition of CHLAMY to SunTag-TRBIP1-MQ1v eliminates genome-wide off-target methylation

The observation that SunTag-NOVA_*FWA*-gRNA robustly induced healthy early-flowering plants in the *fwa* background (Fig. 3A) suggested that it likely was inducing less off-target genome wide DNA methylation as compared to SunTag-TRBIP1-MQ1v_*FWA*-gRNA. To test this, we used whole genome bisulfite sequencing (WGBS) and found that the addition of CHLAMY virtually eliminated the widespread off-target methylation, and also eliminated the methylation of the chloroplast genome (Fig. 3B and Supplementary Fig. 3A,B). At the *FWA* locus, SunTag-NOVA_*FWA*-gRNA promoted highly specific CG methylation within the promoter, with markedly less non-specific spreading into adjacent regions compared with both SunTag-MQ1v_*FWA*-gRNA and SunTag-TRBIP1-MQ1v_*FWA*-gRNA (Fig. 1F,3C). Together, these results demonstrate that SunTag-NOVA serves as an effective epigenetic editing tool capable of depositing methylation at a defined genomic target without inducing genome-wide off-target methylation.

**Fig. 3.**
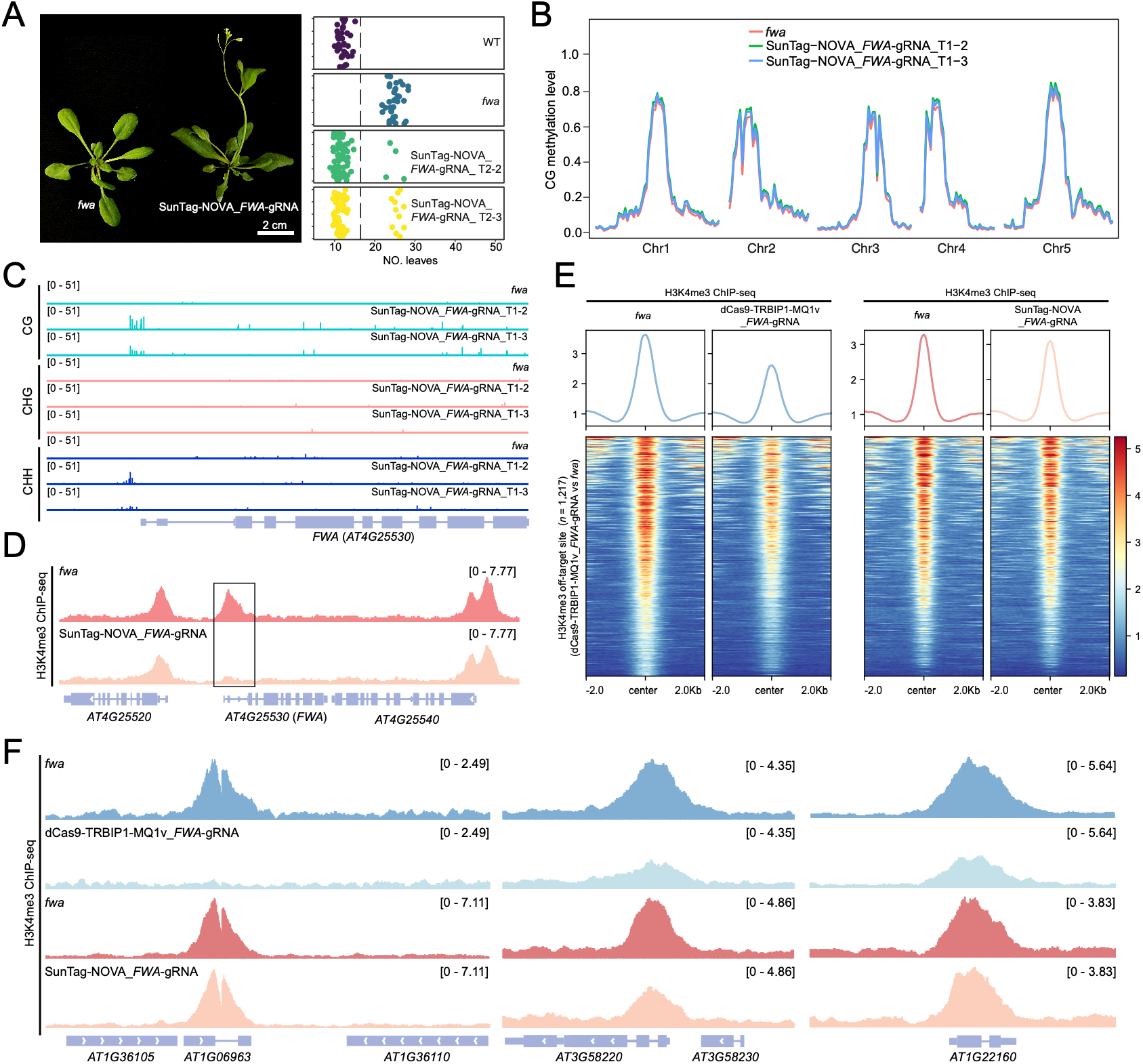
SunTag-NOVA heritably induces DNA methylation at *FWA* without genome-wide off-target effects. (*A*) Early-flowering phenotype of SunTag-NOVA_*FWA*-gRNA T1 plants and leaf numbers of SunTag-NOVA_*FWA*-gRNA T2 lines in the *fwa* background. (*B*) Genome-wide CG methylation levels in SunTag-NOVA_*FWA*-gRNA T1 plants in the *fwa* background. (*C*) Genome browser view showing DNA methylation patterns across the *FWA* promoter in *fwa* and SunTag-NOVA T1 plants. (*D*) Genome-browser tracks showing H3K4me3 ChIP-seq signals in *fwa* and SunTag-NOVA_*FWA*-gRNA T2 plants in the *fwa* background. Black box marks the target *FWA* region, where H3K4me3 enrichment was removed following SunTag-NOVA targeting. (*E*) Metaplots and heat maps comparing H3K4me3 ChIP-seq signals between *fwa* and dCas9-TRBIP1-MQ1_FWA-gRNA T2 plants, and between *fwa* and SunTag-NOVA_FWA-gRNA T2 plants, across H3K4me3 off-target sites identified in dCas9-TRBIP1-MQ1_*FWA*-gRNA plants (signalValue > 1.5 relative to *fwa*). (*F*) Genome-browser tracks showing H3K4me3 ChIP-seq signals (RPGC) at representative off-target sites exhibiting H3K4me3 demethylation in dCas9-TRBIP1-MQ1v_*FWA*-gRNA T2 plants.

Consistent with our previous observation of dCas9-TRBIP1-MQ1v-mediated H3K4me3 removal at the *FWA* locus^7^, chromatin immunoprecipitation followed by sequencing (ChIP-seq) showed a pronounced decrease in H3K4me3 enrichment at the *FWA* promoter in SunTag-NOVA_*FWA*-gRNA plants compared with the *fwa* epiallele (Fig. 3D). In contrast to the SunTag-NOVA system, re-analysis of our previous dCas9-TRBIP1-MQ1v_*FWA*-gRNA ChIP-seq data identified 1,217 genome-wide off-target sites with reduced H3K4me3^7^, whereas these off-target H3K4me3 demethylation events were largely eliminated in SunTag-NOVA_*FWA*-gRNA plants, demonstrating that incorporation of CHLAMY into the SunTag system markedly improves editing specificity in part by suppressing widespread off-target H3K4me3 demethylation (Fig. 3E,F).

### SunTag-NOVA enables targeted DNA methylation editing at additional loci

To evaluate the general applicability of the SunTag-NOVA system, we targeted the promoter regions of *FLOWERING LOCUS T* (*FT*) and *TOO MANY MOUTHS* (*TMM*) in the WT background. WGBS revealed efficient targeted DNA methylation at the *FT* locus in T1 transgenic plants. Although the gRNA targeted a specific promoter region, DNA methylation extended beyond the target site, with the highest methylation detected approximately 1.5 kb downstream of the gRNA-binding region (Fig. 4A and Supplementary Fig. 4). This expanded methylation pattern is consistent with the multivalent nature of the SunTag-NOVA system, which may facilitate higher-order molecular clustering of the effectors through both the SunTag scaffold and the CHLAMY module. We also observed DNA methylation at the targeted *TMM* promoter (Fig. 4A and Supplementary Fig. 4). Quantitative RT-PCR showed that the expression of both *FT* and *TMM* was significantly reduced in independent SunTag-NOVA transgenic lines (Fig. 4B). Consistent with the established biological functions of these genes, repression of *FT* resulted in a late-flowering phenotype, whereas repression of *TMM* caused stomatal clustering (Fig. 4C,D)^16, 17^. Importantly, genome-wide and chloroplast DNA methylation profiles of the SunTag-NOVA transgenic plants were nearly indistinguishable from those of WT plants, indicating that targeted DNA methylation was achieved with minimal genome-wide off-target effects (Fig. 4E and Supplementary Fig. 5A,B). Collectively, these results demonstrate that SunTag-NOVA is an efficient and highly specific targeted DNA methylation editing tool that can be applied to endogenous loci beyond *FWA*.

**Fig. 4.**
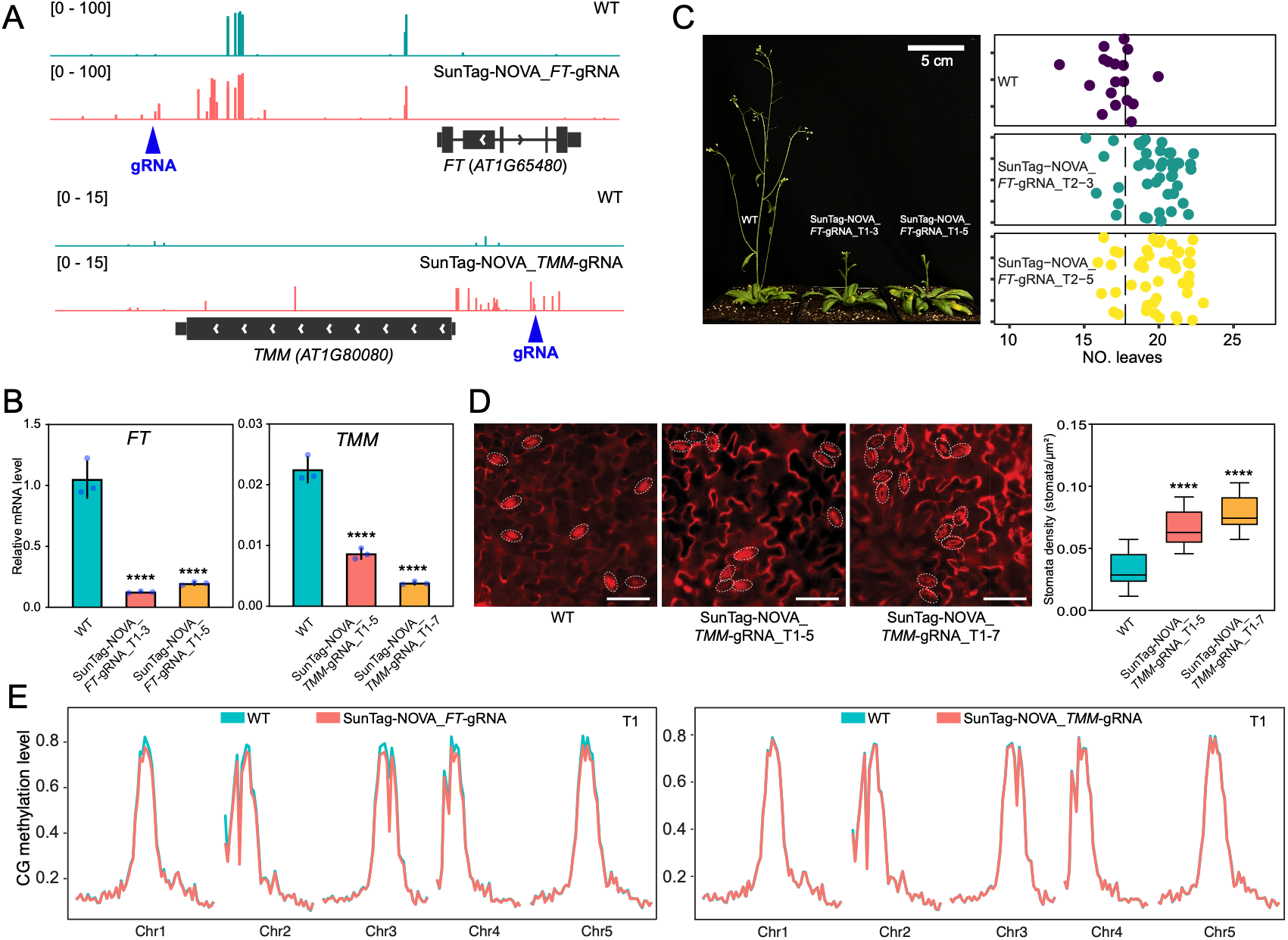
Validation of SunTag-NOVA at endogenous *FT* and *TMM* loci. heritably induces DNA methylation without genome-wide off-target effects. (*A*) CG methylation profiles at the *FT* and *TMM* loci in WT and representative SunTag-NOVA T1 transgenic plants. Blue arrowheads indicate the gRNA target sites. (*B*) Relative expression levels of *FT* and *TMM* in WT and independent SunTag-NOVA T1 transgenic lines. Expression levels were normalized to the internal reference gene *IPP2*. Data are presented as mean ± SD (*n* = 3). Asterisks indicate significant difference by two-tailed Student’s *t*-test (\*\*\*\**P* < 0.0001). (*C*) Representative flowering phenotypes of WT and SunTag-NOVA T1 transgenic plants targeting *FT*. Leaf number is shown as a quantitative measurement of flowering time. Each dot represents one individual plant. (*D*) Representative stereomicroscope images of propidium iodide (PI)-stained leaves from WT and SunTag-NOVA T1 transgenic plants targeting *TMM*, showing stomatal clustering in the transgenic lines. The right panel shows the quantification of stomatal clusters. Dashed circles indicate the positions of stomata. Scale bars, 50 μm. Data are presented as mean ± SD (*n* = 10). Asterisks indicate significant difference by two-tailed Student’s *t*-test (\*\*\*\**P* < 0.0001). (*E*) Genome-wide CG methylation levels in WT and SunTag-NOVA T1 transgenic plants. Metaplots show average DNA methylation levels across chromosomes, demonstrating that targeted methylation editing by SunTag-NOVA does not induce detectable genome-wide DNA methylation changes.

## Discussion

We previously developed the SunTag based epigenome-editing tools SunTag-NtDRMcd and SunTag-MQ1v^4, 6^, which have been utilized in multiple species^7, 18-20^. SunTag-NtDRMcd preferentially induces CHH methylation, consistent with the role of NtDRM in *de novo* DNA methylation through the RdDM pathway. However, the system also induces genome wide and chloroplast DNA methylation. This off-target methylation persists even after removal of the SV40-type nuclear localization signal (NLS), which was deleted to avoid excessive nuclear accumulation of NtDRMcd^6, 21^. By contrast, SunTag-MQ1v exhibits only moderate editing efficiency and still causes weak genome-wide off-target methylation when targeting the *FWA* locus, a locus that is particularly amenable to and widely used for plant epigenome studies^4^. More recently, He et al. developed a series of DNA methylation-editing tools that offer various trade-offs between high editing efficiency and genome-wide off-target methylation^22^.

To overcome the limitations of these systems, we engineered the SunTag-MQ1v system by incorporating the *Chlamydomonas reinhardtii* ACD protein CHLAMY and the H3K4me3-demethylase-associated protein TRBIP1, thereby generating SunTag-CHLAMY-TRBIP1-MQ1v (SunTag-NOVA). Direct fusion of CHLAMY and TRBIP1 to the SunTag-MQ1v platform substantially increased targeted DNA methylation efficiency while also eliminating detectable genome-wide off-target methylation. H3K4me3 is typically enriched at the promoters of actively transcribed genes and is generally antagonistic to DNA^7, 23, 24^. This antagonism may help explain why the targeted installation of DNA methylation at loci other than *FWA* often is unsuccessful. We recently found that TRBIP1 recruits the H3K4me3 demethylase JMJ14 through TRB proteins, thereby promoting the removal of H3K4me3 at target loci^7, 25^. Consistent with this mechanism, targeting TRBIP1 to *FWA* in the *fwa* epiallele using the SunTag system removed H3K4me3 levels at the *FWA* promoter.

ACD proteins form stable dimers that can further assemble into large, dynamic oligomers^13^. We propose that incorporating CHLAMY into the SunTag system improves editing performance through two complementary mechanisms. First, ACD-mediated oligomerization may increase the local concentration of SunTag complexes at dCas9-targeted loci. Second, CHLAMY may sequester unbound effector complexes into large oligomeric assemblies, reducing their availability for recruitment to nonspecific DNA and thereby limiting genome-wide off-target DNA methylation. Together, the incorporation of TRBIP1 and CHLAMY into SunTag-MQ1v enhances both the efficiency and specificity of targeted DNA. Overall, our engineering strategy upgraded SunTag-MQ1v to SunTag-CHLAMY-TRBIP1-MQ1v (SunTag-NOVA), a platform with high targeted DNA methylation efficiency, specificity, and potential applicability across diverse loci. SunTag-NOVA thus provides a promising additional strategy for the epigenetic manipulation of otherwise difficult-to-edit genes across diverse plant species. We also propose that incorporating ACDs such as CHLAMY into other CRISPR based genome editing systems may increase specificity in other eukaryotic organisms.

## Materials and Methods

### Plant Materials and Growth Conditions

In this study, *Arabidopsis thaliana* ecotype Columbia-0 (wild-type, WT) and *fwa* mutant were grown in soil under long-day greenhouse conditions (16 h light / 8 h dark). Transgenic plants were generated using the Agrobacterial-mediated floral dipping method, and at least 24 independent T1 transgenic lines were obtained for each construct.

### Plasmid Construction

The construction of SunTag-MQ1v with two combinations of guides targeting *FWA* promoter was previously described^4, 6^. For the constructions of SunTag-TRBIP1-MQ1v and SunTag-CHLAMY-TRBIP1-MQ1v with two same combinations of guides targeting *FWA* promoter, the coding sequences of MQ1v, TRBIP1, and CHLAMY were amplified and ligated using overlapping PCR, then cloned to the SunTag vector that digested by the *BsiWI* restriction enzyme (Catalog# R3553L, NEB) via In-Fusion cloning method (Catalog# 639650, Takara), and the TRBIP1 and MQ1v were directly fused, while CHLAMY and TRBIP1 were ligated with the Xten-linker. For the constructions of SunTag-CHLAMY-TRBIP1-MQ1v (SunTag-NOVA) that targeted to *FT* and *TMM*, one guide with gRNA scaffold sequence for each gene were synthesized (with 20-50 bp overlapped sequence), respectively, and then we used infusion to the *MauBI* (Catalog# ER2081, Thermo Scientific) digested SunTag-CHLAMY-TRBIP1-MQ1v no guide (SunTag-NOVA no guide) vector. For SunTag-ACD constructs, coding sequences of ACDs from different species were cloned into the *BsiWI*-digested SunTag vector using the In-Fusion cloning method. For the yeast two hybrid constructions of pGBKT7-CHLAMY/ACD and pGADT7-CHLAMY/ACD, the CHLAMY or ACD fragments were amplified and infusion to destination vectors pGBKT7 or pGADT7 with *NdeI* (Catalog# R0111S, NEB) and *EcoRI* (Catalog# R0101S, NEB) digested. The yeast strain AH109 was used for yeast two hybrid assay and the yeast transformation was conducted using the Frozen-EZ Yeast Transformation II TM Kit (Catalog# T2001, Zymo Research) following the manufacturer’s protocol and modified as previously^26^. Guide RNAs are listed in Supplementary Table 2.

### McrBC-qPCR and RT-qPCR

DNA methylation at the *FWA* locus was quantified by McrBC–qPCR. McrBC (NEB, M0272L), a methylation-dependent restriction endonuclease that cleaves DNA containing 5-methylcytosine, was used for digestion. Genomic DNA was extracted from mature leaves using the CTAB method. Equal amounts of DNA were incubated with McrBC or water (mock control) at 37 °C for 4 h, followed by enzyme inactivation at 65 °C for 20 min. Digested DNA was analyzed by qPCR using *FWA*-specific primers. Lower relative McrBC–qPCR values indicate higher levels of DNA methylation. *FWA* transcript levels were measured by quantitative RT– PCR (RT–qPCR). Total RNA was isolated from mature leaves using the Direct-zol RNA MiniPrep Kit (Zymo Research, R2052). One microgram of total RNA was reverse-transcribed using SuperScript IV VILO Master Mix (Invitrogen, 11756500). qPCR was performed using *FWA*-specific primers with *IPP2* as the internal control. All qPCR reactions were carried out using iQ SYBR Green Supermix (Bio-Rad, 1708882). Primers are listed in Supplementary Table 1.

### BS-PCR Sequencing

Genomic DNA was extracted from mature leaves using the CTAB method. Ten microliters of each DNA sample was subjected to bisulfite conversion using the EpiTect Bisulfite Kit (QIAGEN, 59104) according to the manufacturer’s instructions. Bisulfite-converted DNA was amplified by PCR using PfuTurbo Cx DNA polymerase (Agilent, 600410) to generate amplicons spanning three regions of the *FWA* promoter: region 1 (chr4: 13,038,143–13,038,272), region 2 (chr4: 13,038,356–13,038,499), and region 3 (chr4: 13,038,568–13,038,695). PCR products were purified using AMPure beads (Beckman Coulter, A63881) and subsequently used for library preparation with the KAPA DNA HyperPrep Kit (Roche, KK8502) and TruSeq DNA UD indexes (Illumina). Libraries were sequenced on an Illumina iSeq 100 platform. BS-PCR primers are listed in Supplementary Table 1.

### Whole-genome Bisulfite Sequencing (WGBS)

Genomic DNA was extracted from mature leaves using the DNeasy Plant Mini Kit (QIAGEN, 69106). A total of 100 ng of genomic DNA was used for library preparation. DNA was fragmented at 4 °C for 2 min using a Covaris S2 instrument, followed by end repair and adapter ligation according to the KAPA DNA HyperPrep Kit protocol (Roche, KK8502). 1 μL EM-seq Adapter (E7165A) from NEBNext Enzymatic Methyl-seq v2 Kit (E8015S) were used for ligation. Adapter-ligated DNA was purified using AMPure beads (Beckman Coulter, A63881) and subsequently subjected to bisulfite conversion using the EpiTect Bisulfite Kit (QIAGEN, 59104). Libraries were amplified using 1 μL UDI primers from NEBNext LV Unique Dual Index Primers Set 5 (E3408S) and sequenced on an Illumina NovaSeq 6000 platform. WGBS data were analyzed following a previously described pipeline^27^, and methylation profiles were visualized using ViewBS software (v0.1.11)^28, 29^.

### Chromatin Immunoprecipitation Sequencing (ChIP-seq)

ChIP-seq was performed using approximately 2 g of mature leaf tissue per sample. Tissues were harvested and flash-frozen in liquid nitrogen, then ground to a fine powder and resuspended in 25 mL nuclear isolation buffer (50 mM HEPES, 1 M sucrose, 5 mM KCl, 5 mM MgCl_2_, 0.6% Triton X-100, 0.4 mM PMSF, 5 mM benzamidine, and protease inhibitor cocktail [Sigma, 11873580001]) supplemented with 1% formaldehyde. Crosslinking was carried out at room temperature for 10 min with gentle agitation and quenched by addition of 1.7 mL of 2 M glycine. Samples were filtered through a single layer of Miracloth (EMD Millipore, 475855-1R) and centrifuged at 2,880 × g for 20 min at 4 °C. Pellets were sequentially resuspended in extraction buffer 2 (0.25 M sucrose, 10 mM Tris-HCl pH 8.0, 10 mM MgCl_2_, 1% Triton X-100, 5 mM β-mercaptoethanol, 0.1 mM PMSF, 5 mM benzamidine, and protease inhibitors) and extraction buffer 3 (1.7 M sucrose, 10 mM Tris-HCl pH 8.0, 2 mM MgCl_2_, 0.15% Triton X-100, 5 mM β-mercaptoethanol, 0.1 mM PMSF, 5 mM benzamidine, and protease inhibitors), with centrifugation at 12,000 × g at 4 °C for 10 min and 1 h, respectively. The final nuclear pellet was resuspended in 400 μL lysis buffer (50 mM Tris-HCl pH 8.0, 10 mM EDTA, 1% SDS, 0.1 mM PMSF, 5 mM benzamidine, and protease inhibitors) and diluted with 1.7 mL ChIP dilution buffer (1.1% Triton X-100, 1.2 mM EDTA, 16.7 mM Tris-HCl pH 8.0, 167 mM NaCl, 0.1 mM PMSF, and 5 mM benzamidine). Chromatin was sheared using a Bioruptor Plus (Diagenode, B01020001) for 14 cycles (30 s on / 30 s off). Lysates were clarified by centrifugation at maximum speed at 4 °C for 10 min (twice), and supernatants were incubated with 6 μL anti-H3K4me3 antibody (Abcam, ab213224) at 4 °C overnight. 25 μL Protein A and 25 μL Protein G magnetic Dynabeads (Invitrogen, 10002D and 10004D) were added and incubated for 2 h at 4 °C. Beads were washed sequentially with low-salt buffer (twice), high-salt buffer, LiCl buffer, and TE buffer, each for 5 min at 4 °C. Chromatin was eluted with elution buffer (1% SDS, 10 mM EDTA, 0.1 M NaHCO_3_) at 65 °C for 30 min, followed by reverse crosslinking at 65 °C overnight. Samples were treated with proteinase K at 45 °C for 4 h, and DNA was purified by phenol–chloroform extraction using Phase Lock Gel (VWR, 2302820) and ethanol precipitation with GlycoBlue (Invitrogen, AM9516). DNA pellets were washed with 70% ethanol and eluted in nuclease-free water. ChIP DNA was used for library preparation with the KAPA DNA HyperPrep Kit (Roche, KK8502) according to the manufacturer’s instructions and sequenced on an Illumina NovaSeq X Plus platform.

### Confocal Microscopy

All confocal microscopy experiments were performed using an LSM 980 confocal microscope (Zeiss). For live-cell imaging, root tips from 2-week-old seedlings grown on half-strength Murashige and Skoog (½ MS) medium at room temperature were transferred with forceps onto 1-mm-thick glass slides (Fisher Scientific, 12-550-08) containing deionized water prior to imaging.

### AlphaFold Prediction

Full-length CHLAMY protein and the ACD domain were predicted using the AlphaFold3 server (https://alphafoldserver.com)^30^. The highest-confidence predicted model was selected for subsequent structural visualization and analysis using ChimeraX (v1.10.1).

## Data, Materials, and Software Avalability

The high-throughput sequencing data generated in this paper have been deposited in the Gene Expression Omnibus (GEO) database (Need to be uploaded). Source data are provided with this paper.

## Acknowledgment

We thank all members of the Jacobsen Lab for their helpful insights and valuable suggestions. We also thank Mahnaz Akhavan for support with NGS at the UCLA Broad Stem Cell Research Center BioSequencing Core Facility. This work was supported by Bill and Melinda Gates Foundation (OPP1125410) and W. M Keck Foundation grants to S.E.J., and NSF GRFP (DGE-2034835, DGE-2444110) to T.J.B. S.E.J. is an Investigator of the Howard Hughes Medical Institute.

## Author contributions

Y.H. and S.E.J. designed research; Y.H., M.W., T.J.B., B.A.B., E. L., J. S., N.A., B.X., A.N., and S.F. performed research; Y.H., S.W., and Y.Z analyzed data; and Y.H. and S.E.J. wrote the paper.

