## Supplementary data for "Chromatin-Assisted Targeting Enables Precise DNA Methylation Editing in Plants"

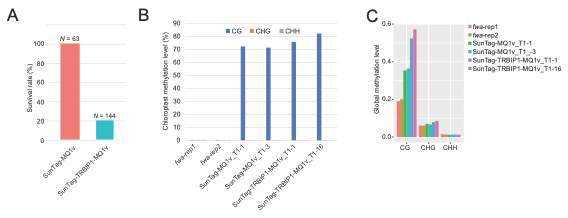

**Supplementary Fig. 1. TRBIP1 within SunTag-MQ1v framework causes lethality and off-target methylation.** (A) Survival rate comparison of T1 plants expressing SunTag-MQ1v or SunTag-TRBIP1-MQ1v in the *fwa* background. (B) Chloroplast methylation indicating off-target methylation. (C) Global methylation level comparison of T1 plants expressing SunTag-MQ1v or SunTag-TRBIP1-MQ1v in the *fwa* background.

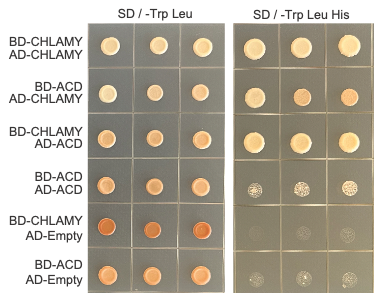

**Supplementary Fig. 2. Y2H validation of self-interaction of CHLAMY to form dimer**

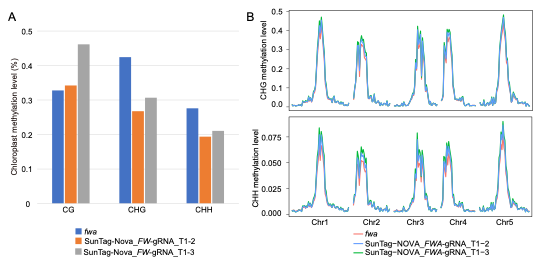

**Supplementary Fig. 3. Minimal off-target DNA methylation in SunTag-NOVA_*FWA*-gRNA plants.** (A) Chloroplast DNA methylation levels in SunTag-NOVA_*FWA*-gRNA T1 plants. (B) Genome-wide CHG and CHH methylation levels in SunTag-NOVA_FWA-gRNA T1 plants in *fwa*.

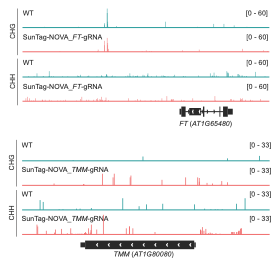

**Supplementary Fig. 4. CHG and CHH methylation profiles at the FT and TMM loci in WT and representative SunTag-NOVA T1 plants.**

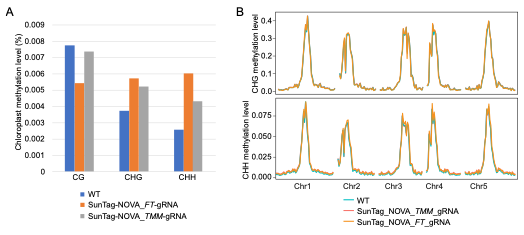

**Supplementary Fig. 5. Minimal off-target DNA methylation in SunTag-NOVA_*FT*-gRNA and SunTag-NOVA_*TMM*-gRNA plants.** (A) Chloroplast DNA methylation levels in SunTag-NOVA_*FT*-gRNA and SunTag-NOVA_*TMM*-gRNA T1 plants in WT. (B) Genome-wide CHG and CHH methylation levels in SunTag-NOVA_*FT*-gRNA and SunTag-NOVA_*TMM*-gRNA T1 plants in the WT backgound.

**Supplementary Table 1. Primers used in this study**

| **Name** | **Sequence** |
| --- | --- |
| IPP2_qPCR_F | GTATGAGTTGCTTCTCCAGCAAAG |
| IPP2_qPCR_R | GAGGATGGCTGCAACAAGTGT |
| FWA_qPCR_F | TTAGATCCAAAGGAGTATCAAAG |
| FWA_qPCR_R | CTTTGGTACCAGCGGAGA |
| FT_qPCR_F | GGAGACGTTCTTGATCCGTTTA |
| FT_qPCR_R | CAATGGAGATATTCTCGGAGGTG |
| TMM_qPCR_F | CTGGTCCTGTCAGACTGTTATC |
| TMM_qPCR_R | CCTTCACCTAGAGGGCAATAAT |
| FWA_McrBC-F | TTGGGTTTAGTGTTTACTTG |
| FWA_McrBC-R | GAATGTTGAATGGGATAAGGTA |
| FWA_BS-PCR_1F | TCATATAAAAAAAAAATTAAATTTCATTTCACAATAACCATT |
| FWA_BS-PCR_1R | GTATGGGYTTYGATAAAGAATATATGAGATTYT |
| FWA_BS-PCR_2F | CTCATATATACCTTATCCCATTCAACATTCATA |
| FWA_BS-PCR_2R | AAGATYTGATATTTGGYTGGAAAAAAYAATAATAAT |
| FWA_BS-PCR_3F | CRCTCTTTATCCCATTCAACATTCATAC |
| FWA_BS-PCR_3R | TTTGGTTGAAAAAAATAATAAAAATTTGATTGTYAGTAT |

**Supplementary Table 2. Guide RNA used in this study**

| **Guide RNA** | **Sequence** |
| --- | --- |
| FWA_gRNA_4 | acggaaagatgtatgggctt |
| FWA_gRNA_7 | aaaactaggccatccatgga |
| FT_gRNA | ctcttcgaattacattcgta |
| TMM_gRNA | aatctgaataaacggacgcg |
